# A deep dive into the *FALSIFLORA* contribution to tomato reproductive development through a comprehensive allelic series analysis

**DOI:** 10.1101/2024.05.21.595162

**Authors:** Abraham S. Quevedo-Colmena, Wim H. Vriezen, Pieter G.A. Wesselink, José M. Pérez, Benito Pineda, Begoña García-Sogo, Trinidad Angosto, Vicente Moreno, Fernando J. Yuste-Lisbona, Rafael Lozano

## Abstract

Plants undergo continuous growth and development thanks to meristems, specialized groups of pluripotent stem cells that remain in their undifferentiated state throughout the plant’s life cycle. Meristem transition from the vegetative to the reproductive phase heavily influences plant reproductive success and agricultural productivity. In tomato (*Solanum lycopersicum* L.), *FALSIFLORA* (*FA*), the orthologue of the Arabidopsis *LEAFY* gene, promotes floral transition by specifying floral meristem identity and regulating the expression of genes responsible for floral organ identity and development. In this study, the allelic series of *FA* has been expanded by combining the screening of a tomato EMS mutant collection with overexpression, gene silencing and CRISPR/Cas9 genome editing approaches, aimed to deepen the understanding of the functional role of *FA* during tomato reproductive development. The phenotypic and molecular characterization of the *FA* allelic series revealed the multifaceted role of *FA* acting in both early and late stages of floral ontogeny. Thus, in addition to promoting floral transition and specifying floral meristem identity, *FA* also plays a role in inflorescence meristem maturation and termination, thereby regulating the inflorescence architecture. Furthermore, *FA* exerts regulatory control over the expression of the tomato homologs of *AGAMOUS* (*TOMATO AGAMOUS1*, *TAG1*) and *WUSCHEL* (*SlWUS*), underscoring its function in promoting carpel development and suppressing floral stem cell activity, thereby establishing floral determinacy. In conclusion, our findings demonstrate the potential of employing mutant allelic series as powerful tools for elucidating gene functions and deciphering the intricate molecular basis underlying biological processes.

**Plain Language Summary:** Our research aims to gain a deeper understanding of the role of *FALSIFLORA* in tomato reproductive development. The *FALSIFLORA* allelic spectrum was expanded through mutant screening, along with overexpression, silencing, and CRISPR/Cas9 editing approaches. Our findings reveal that *FALSIFLORA* has a dual function in early and late floral development stages. In addition to promoting floral transition and specifying floral meristem identity, *FA* regulates the expression of tomato genes homologous to *WUSCHEL* and *AGAMOUS*, which play an essential role in stem cell regulation and carpel development. This study highlights the value of allelic series in elucidating gene functions and intricate biological mechanisms.

## Introduction

The transition from vegetative growth to flowering stands out as a pivotal event in the life cycle of a plant. This transition is of significant importance not only in the reproductive phase, but also in influencing important agronomic traits in crop species. In monopodial plants such as the model species *Arabidopsis thaliana* L. and *Antirrhinum majus* L., the vegetative-to-floral transition initiates with the generation of an inflorescence meristem (IM) from the indeterminate shoot apical meristem. This IM gives rise to a determinate floral meristem (FM), which then differentiates into a flower, while a new IM is initiated at its flank. In contrast, in plants with sympodial growth such as tomato (*Solanum lycopersicum* L.), the shoot apex is determined and converted into an IM. Vegetative development continues from a specialized axillary meristem called the sympodial meristem (SYM), which leads to the formation of a variable number of leaves and a new terminal inflorescence. Hence, the sympodial pattern is characterized by a regular alternation of vegetative and reproductive phases. In tomato, after the induction of floral development, the formed IM produces an FM, and a new lateral IM arises adjacently. Thus, each IM originates another IM while maturating to FM, thereby establishing a sequential growth dynamic that results in the production of an inflorescence organized in a zig-zag pattern (reviewed in Lozano et al. 2009; Thouet et al. 2012).

The transition to flowering involves the activation of specific genes that control the timing and formation of floral buds in response to endogenous and environmental cues. In Arabidopsis, the *LEAFY* (*LFY*) gene plays a central role in promoting FM identity and initiating the floral organ formation (Huala and Sussex 1992; Weigel et al. 1992). The *LFY* gene encodes a pioneer plant-specific transcription factor capable of binding to nucleosomes in closed chromatin regions, opening local chromatin by displacing H1 linker histones and recruiting the SWItch Sucrose Non-Fermentable (SWI/SNF) chromatin remodeling complex. Thus, by reprogramming the epigenetic landscape, LFY leads to the establishment of gene expression competence, directly or indirectly enabling other transcription factors to bind to their target genes involved in floral fate (Jin et al. 2021; Lai et al. 2021; Yamaguchi 2021). Structural studies have shown that LFY is a helix-turn-helix transcription factor that contains a sterile alpha motif (SAM) oligomerization N-terminal domain, which is essential for LFY floral function and required to access regions with low-affinity binding sites and closed chromatin (Sayou et al. 2016). In addition, LFY possesses a C-terminal DNA-binding domain (DBD), which is a helix-turn-helix fold that self-dimerizes on DNA and recognizes semi-palindromic 19-bp *cis*-elements (Moyroud et al. 2011; Winter et al. 2011; Sayou et al. 2016).

The function of the *LFY* gene is evolutionarily conserved in the distantly related species *A. majus*, where *FLORICAULA* (*FLO*), the *LFY* orthologue, controls FM identity (Carpenter and Coen 1990; Coen et al. 1990). In tomato, *FALSIFLORA* (*FA*) is orthologous to *LFY* and *FLO*, and plays a role in promoting floral transition and determining FM identity (Molinero-Rosales et al. 1999). Thus, defects reported for the *fa* null allele include delayed flowering, altered sympodial development, and a distinctive phenotype characterized by the conversion of flowers into indeterminate vegetative shoots, resulting in an abnormal inflorescence architecture (Allen and Sussex 1996; Molinero-Rosales et al. 1999). In addition, hypomorphic alleles of *FA* have also been identified in tomato, known as *leafy inflorescence* (*fa^lfi^*) and *pistillate* (renamed as *fa^pi^*). The *fa^lfi^* mutant does not develop flowers, but some leafy organs can transform into fleshy carpelloid structures capable of ripening to a red color (Kato et al. 2005). In contrast, the *fa^pi^* mutant is able to develop flowers, albeit with aberrant organs, and its main feature is a pronounced modification or apparent absence of the staminal cone (Olimpieri and Mazzucato 2008). Consequently, the *fa^pi^*mutant phenotype reveals that *FA* also regulates floral organ identity genes, consistent with the role previously described for *LFY* as a transcriptional activator of genes involved in the ABC model (Huala and Sussex 1992; Weigel and Meyerowitz 1993). LFY binds to the regulatory elements of the C-class MADS-box gene *AGAMOUS* (*AG*) and cooperates with the stem cell regulator WUSCHEL (WUS), a homeobox transcription factor, to activate *AG* expression (Lenhard et al. 2001; Lohmann et al. 2001). In turn, the *AG* homeotic gene has a dual role in specifying organ fate and limiting stem cell proliferation in flowers through transcriptional repression of *WUS*, a process known as floral determinacy (reviewed in Sun and Ito 2015). As a result, *ag* mutants exhibit stem cell maintenance phenotypes, leading to the formation of flowers within flowers, and homeotic transformation of stamens to petals (Bowman et al. 1989). In tomato, although *fa^pi^*flowers occasionally exhibit indeterminacy and the development of sepal-like structures inside the ovary (Olimpieri and Mazzucato 2008), the role of *FA* in floral determinacy has so far not been addressed.

Valuable understanding of the functional role of a gene, its molecular mechanisms, and its contribution to biological processes can be obtained by systematically studying an allelic series. In this study, novel alleles of the *FA* locus have been identified and generated by combining the screening of a tomato ethyl methanesulfonate (EMS) mutant collection with overexpression, gene silencing and CRISPR/Cas9 genome editing methods. A comprehensive phenotypic and molecular characterization of a series of *fa* mutant alleles allowed us to gain insights into the role of *FA* in tomato reproductive development. Our results demonstrate that *FA* functions in both early and late stages of floral development. Thus, *FA* not only promotes floral transition and FM identity, but also plays a role in floral determinacy by regulating both the tomato homologues of *WUS* (*SlWUS*) and *AG* (*TOMATO AGAMOUS1*, *TAG1*). Taken together, our data reveal the potential of a mutant allele series to decipher gene functions and unravel the molecular basis of key biological processes.

## Material and methods

### Plant material and growth conditions

The *fa^bif^* mutant was identified in a screening of a mutant collection obtained by chemical mutagenesis with EMS in *S. lycopersicum* cv. TPAADASU (Nunhems; Gady et al. 2009). Both, the *fa* (Ailsa Craig background, accession LA0854) and *fa^pi^* (San Manzano background, accession 2-137) lines were obtained from the Tomato Genetics Resource Center (http://tgrc.ucdavis.edu/) and were used along with the *fa^bif^* line for the characterization of reproductive development. All experiments were conducted under greenhouse conditions following standard management practices, including regular fertilization.

### Measurements of reproductive traits

To determine potential differences in reproductive phenotypic traits between *fa* mutants and their respective wild-type counterparts, we assessed (i) flowering time, measured as the number of leaves before flowering; (ii) inflorescence architecture, comprised by variations in the number of flowers or leaves for the second, third, and fourth inflorescences per plant; and (iii) floral organ number. The phenotypic traits, measured on 15 plants and 5 flowers of each genotype, were considered significantly different when the Student’s t-test P-values were < 0.05.

### Microscopy analysis

For the study of inflorescence architecture, reproductive meristems from wild-type and mutant genotypes were analyzed using a Leica DMS1000 Stereo microscope. Inflorescence development and floral organ identity were assessed by scanning electron microscopy (SEM) analyses, following the procedure described in Lozano et al. (1998). Samples were fixed in FAEG solution (comprising 10% formaldehyde, 5% acetic acid, 50% absolute ethanol, and 0.72% glutaraldehyde) for 72 hours and then stored in 70% ethanol. Before analysis, samples were dehydrated in increasing ethanol concentrations and dried using a CO_2_ critical point dryer (Bal-Tec CPD 030). Subsequently, the samples were gold-coated using a LEICA EM ACE 200 sputter coater, and visualized in a high-resolution FESEM-Zeiss Sigma 300 VP scanning electron microscope at 10 kV. Furthermore, alterations in floral determinacy were evaluated by light microscopy. Samples were fixed in FAE solution (10% formaldehyde, 5% acetic acid, and 50% absolute ethanol), followed by ethanol dehydration, embedding in paraffin, and sectioning using a Leica RM2035 microtome. Eight-µm thick transverse sections were stained with a 1% toluidine blue solution and examined under a Leica DM6 B microscope.

### Molecular analysis of *fa*, *fa^pi^* and *fa^bif^* alleles

Genomic DNA was extracted from young leaves using the DNAzol® Reagent kit (Invitrogen Life Technologies, San Diego, CA, USA) and quantified using a NanoDrop 2000 spectrophotometer (Thermo Scientific, Wilmington, DE, USA). Based on the deletion/frameshift mutation described in Molinero-Rosales et al. (1999), the *fa* mutant allele was genotyped using a CAPS marker that amplified a 396 bp fragment. Subsequent digestion with *Ssi*I cleaved the 396 bp fragment from the wild-type allele into 195, 120, and 81 bp fragments. However, the *fa* mutation eliminates one of the *Ssi*I recognition sites, leading to the production of 260 and 120 bp fragments. To identify the molecular nature of the *fa^pi^*and *fa^bif^* mutations, the *FA* (*Solyc03g118160*) genomic region was sequenced by Sanger using the ABI PRISM BigDye Terminator Cycle Sequencing kits on an ABI PRISM 3500 Genetic Analyzer (Applied Biosystems, Foster City, CA, USA). The CAPS marker designed for genotyping the *fa^pi^* mutation generated a 191 bp amplicon. Upon treatment with *Taq*I, the mutant allele is cleaved into 164 and 27 bp fragments. Finally, a dCAPS marker was designed to genotype the *fa^bif^* mutation, resulting in a 701 bp fragment. After the *Tru*I treatment, the amplicon derived from the *fa^bif^* mutant allele is cleaved into 399 and 302 bp fragments. Primers used to obtain *FA* coding sequences and genotyping markers are listed in Supplementary Table S1.

### Generation of transgenic lines

To generate the *35S:FA* overexpression construct, the full-length coding sequence of *FA* was amplified using Phusion™ High-Fidelity DNA polymerase (Thermo Fisher Scientific, Waltham, MA, USA) and gene-specific primer pairs (Supplementary Table S2). It was then cloned into the pENTR/D-TOPO vector and subsequently subcloned into the Gateway pGWB402 binary vector (Nakagawa et al. 2007) using the Gateway LR Clonase system (Invitrogen, Karlsruhe, Germany). To generate the RNA interference (RNAi) *FA* construct, a 165-bp fragment from the first exon of *Solyc03g118160* was amplified using the FA_RNAi_F and FA_RNAi_R primers (Supplementary Table S2) and inserted into the pKannibal vector in both sense and antisense orientations (Wesley et al. 2001). The modified pKannibal vector was digested with NotI, and the resulting restriction fragment was then integrated into the pART27 vector (Gleave 1992). The CRISPR/Cas9 constructs for editing both the *FA* coding (*fa^CR-cds^*) and promoter (*fa^CR-pro^*) sequences were generated according to the protocol described by Vazquez-Vilar et al. (2016). Single-guide RNAs (sgRNAs) were designed using the Breaking-Cas software (Oliveros et al. 2016). One sgRNA targeting the *FA* coding region was used to construct the *fa^CR-cds^*vector, while three *fa^CR-pro^* vectors, each containing three sgRNAs, were developed to induce cis-regulatory mutations. These sgRNAs were designed to target the 4-kb promoter region situated immediately upstream of the *FA* coding sequence. The primers used to generate of CRISPR/Cas9 constructs are shown in Supplementary Table S2. Genetic transformation experiments were conducted using *Agrobacterium tumefaciens* (strain LBA4404), according to the methodology described by Ellul et al. (2003). The ploidy levels in transgenic plants were determined by flow cytometry, according to the protocol described by Atares et al. (2011). Transgenic plants were grown under greenhouse conditions, using standard management practices, including regular fertilization. To differentiate lines carrying edited alleles from wild-type lines, *fa^CR-cds^* and *fa^CR-pro^* CRISPR/Cas9 plants were genotyped using primers designed to cover the target recognition regions of the sgRNAs (Supplementary Table S3). PCR products were purified and cloned into the pGEMT vector (Promega, Madison, WI, USA). A minimum of 10 clones per PCR product were then sequenced by the Sanger method to characterize the edited alleles.

### RNA isolation and quantitative real-time PCR analysis

Total RNA was extracted using TRIzol reagent (Invitrogen Life Technologies, San Diego, CA, USA), and genomic DNA contamination was removed by treatment with the DNA-free™ kit (Ambion, Austin, TX, USA). One microgram of RNA was used for cDNA synthesis using a ML-MLV reverse transcriptase (Invitrogen, San Diego, CA, USA) with a mixture of random hexamer and oligo-(dT)18 primers. qRT-PCR analysis was carried out using three biological and two technical replicates. Specific primer pairs (Supplementary Table S4) were used in each qRT-PCR with the SYBR Green PCR Master Mix Kit (Applied Biosystems, Foster City, CA, USA) on the QuantStudio 1 Real-Time PCR System (Applied Biosystems, Foster City, CA, USA). The ΔΔCt method (Winer et al. 1999) was employed to calculate relative gene expression levels, with *UBIQUITIN3* (*Solyc01g056940*) as a housekeeping gene.

## Results

### An allelic series of mutations in *FA* shows variation in flowering time and inflorescence branching

In a mutagenesis experiment designed to identify regulators of the inflorescence development, we identified a recessive mutant named as *branched inflorescences and indeterminate fruits* (*bif*). This mutant produces extremely compound inflorescences bearing leaves and non-fertile flowers with aberrant organs, which develop indeterminate fruits (Fig. 1l, Fig. 3i, and Fig. 4f). As this phenotype is reminiscent of that occasionally described for *fa* hypomorphic alleles, we conducted an allelism test with the *fa* null mutant. In the F1 progeny derived from the cross of a heterozygous *bif* plant with a heterozygous *fa* plant, a 3:1 ratio of wild-type:mutant phenotype was observed (37 wild-type:11 mutant, χ^2^ = 0.11, *P* = 0.74), proving that *bif* is a new allele of the *FA* gene and was therefore renamed *fa^bif^*.

**Fig. 1.**
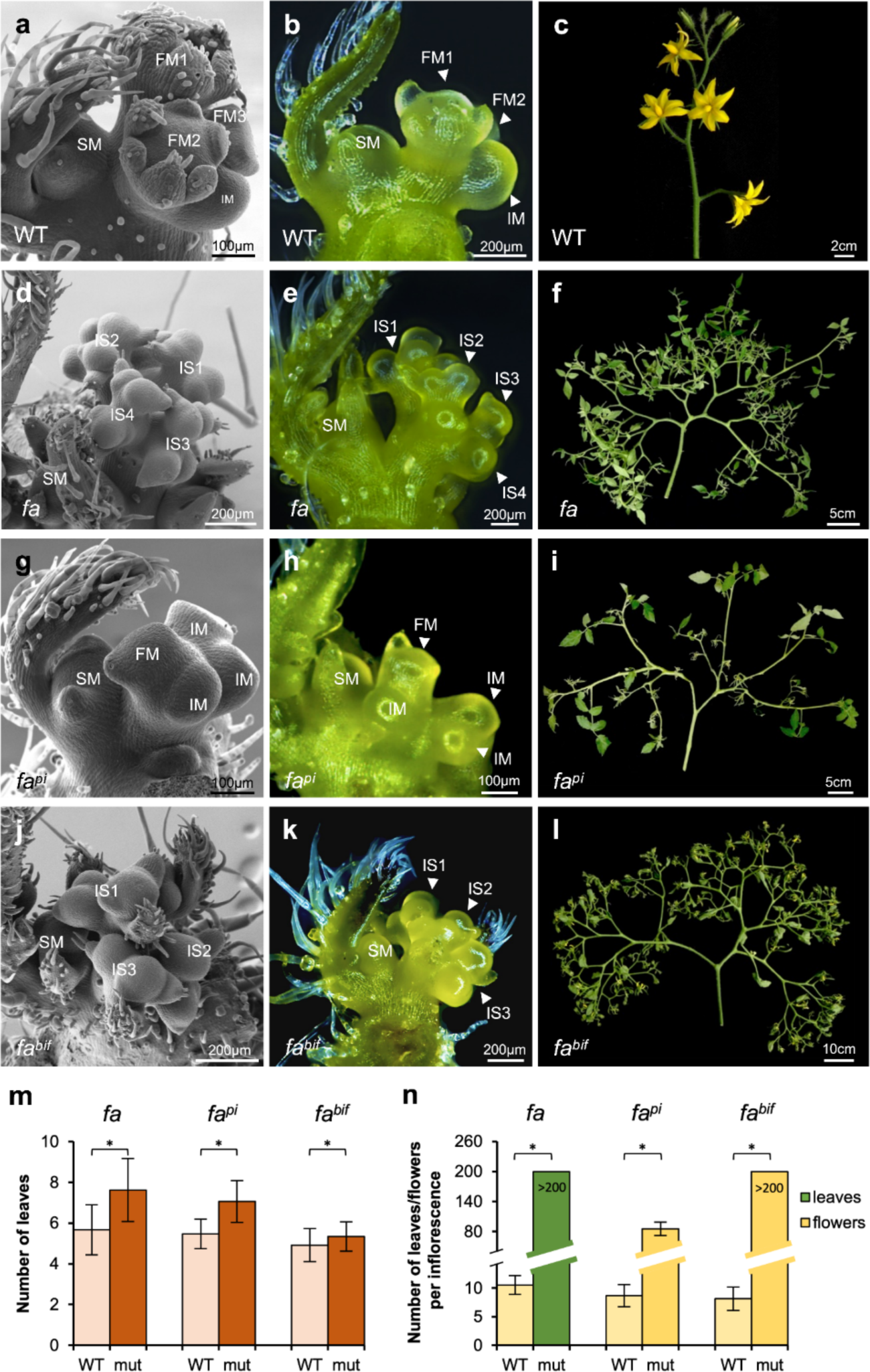
Mutations at the *FA* locus alter flowering time and inflorescence architecture. Scanning electron microscope (SEM) images (a, d, g, j) and stereoscopic images (b, e, h, k) illustrating inflorescence development of wild-type (WT) (a-c), *fa* (d-f), *fa^pi^* (g-i) and *fa^bif^* (j-l) plants. SM, sympodial meristem; IM, inflorescence meristem; FM, floral meristem; IS, inflorescence shoot. Inflorescence architecture of WT (c), *fa* (f), *fa^pi^*(i) and *fa^bif^* (l) plants after 80 days after sowing. Number of leaves before flowering (m) and number of leaves/flowers per inflorescence (n) in WT, *fa*, *fa^pi^* and *fa^bif^*. ns, not significant difference; * significant differences at *P* < 0.05. (Student’s t-test).

**Fig. 2.**
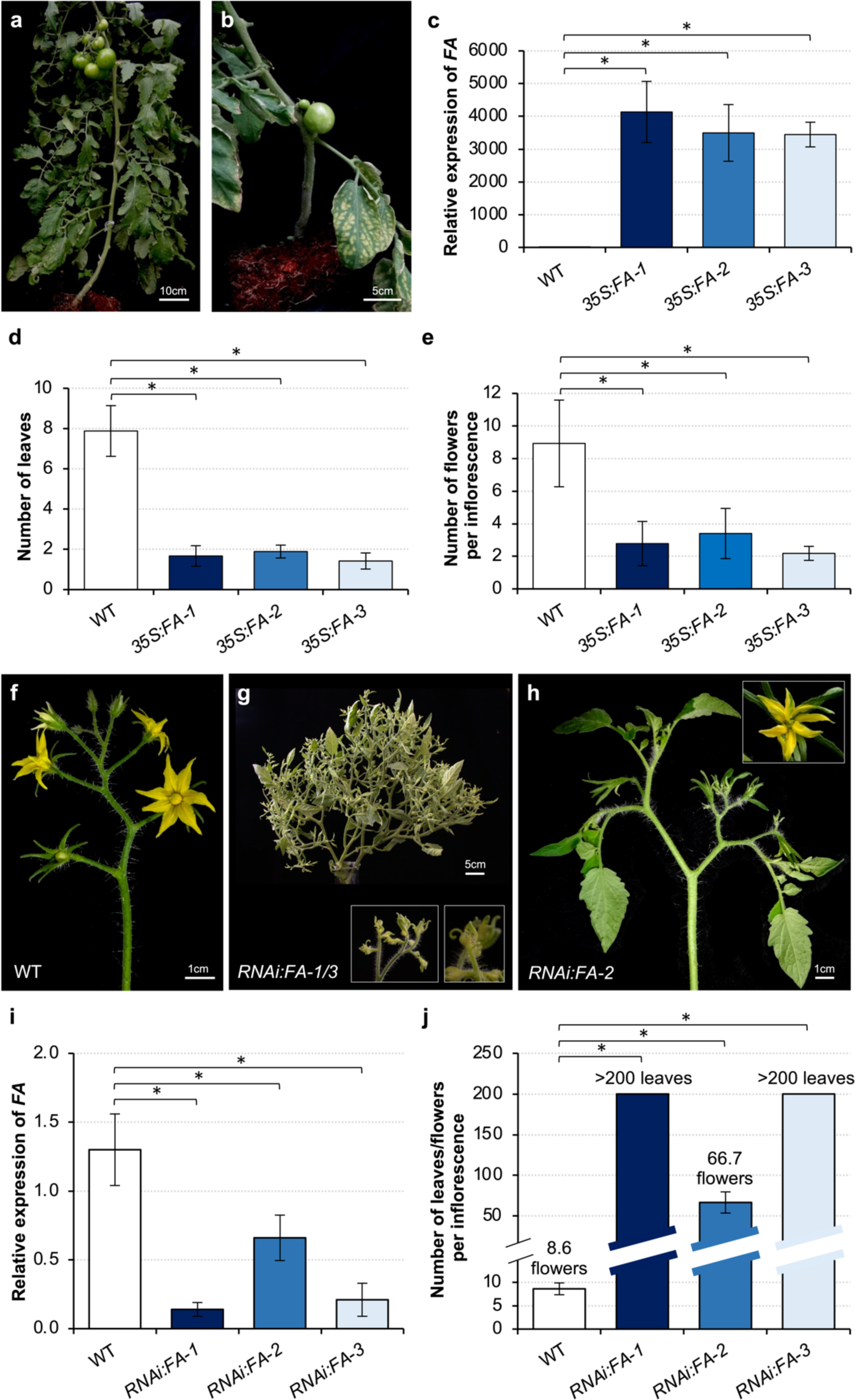
Effects of overexpression and RNA interference (RNAi)-mediated knockdown of the *FA* gene. (a-e) Overexpression of *FA* caused a reduction in flowering time and the number of flowers developed per inflorescence. (a, b) Representative phenotypes of wild-type (WT) and *FA* overexpression (*35S:FA*) plants. (c) Relative *FA* expression in WT and three independent two-generation (TG2) *35S:FA* lines. Number of leaves before flowering (d) and number of flowers per inflorescence (e) in WT and *35S:FA* lines. (f-j) RNAi-mediated knockdown of *FA* promotes the formation of branched inflorescences. (f, g, h) Representative phenotypes of WT and *RNAi:FA* plants. (i) Relative *FA* expression in WT and three independent first-generation (TG1) *RNAi:FA* lines. (j) Number of leaves/flowers per inflorescence in WT and *RNAi:FA* lines. ns, no significant difference; * significant differences at P < 0.05. (Student’s t-test).

**Fig. 3.**
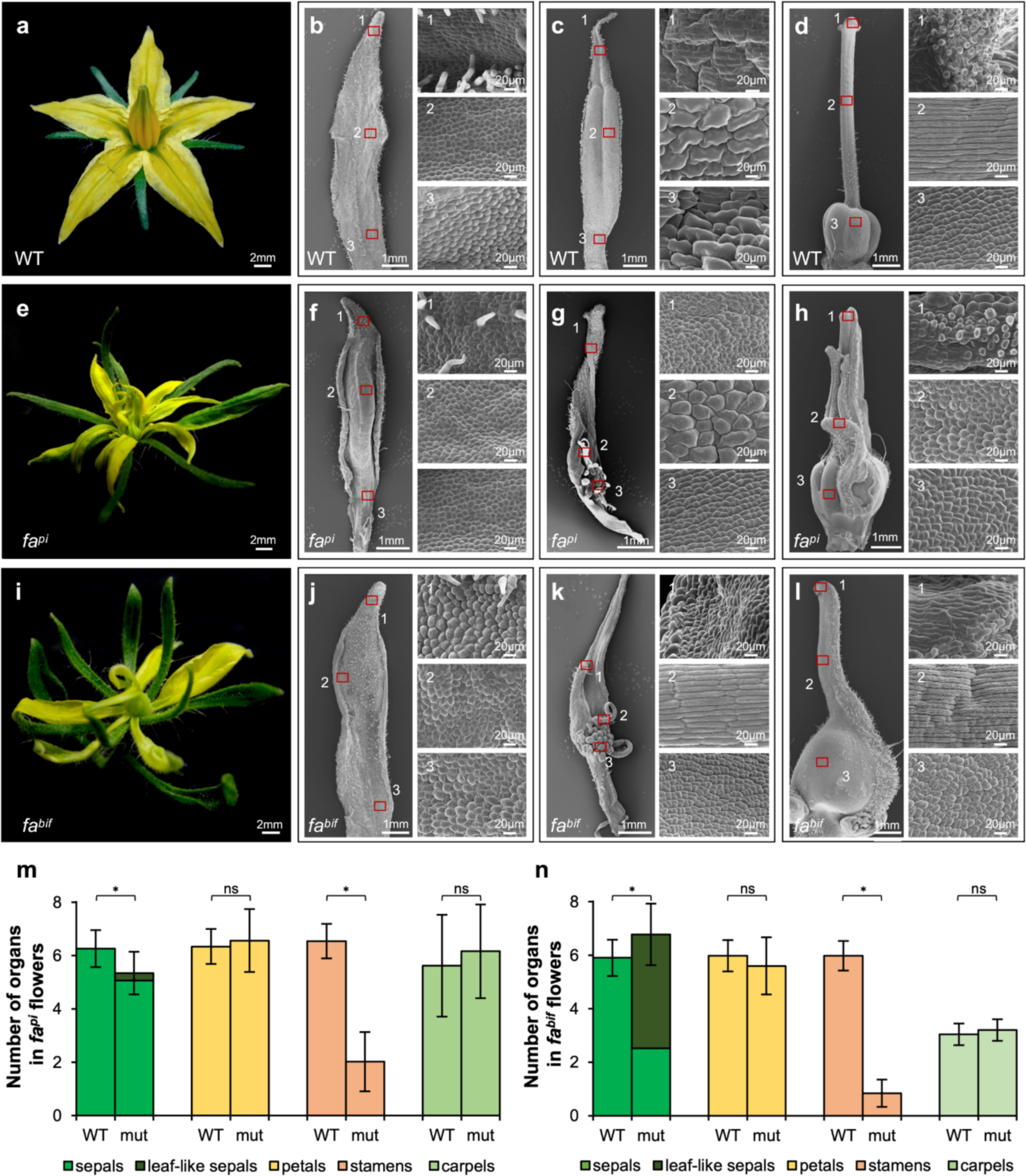
The *fa^pi^* and *fa^bif^* mutants develop flowers with homeotic transformations and a reduced number of floral organs. Representative flowers of wild-type (WT) (a), *fa^pi^* (e), and *fa^bif^* (i) plants. Scanning electron microscope (SEM) images of petals (b,f,j), stamens (c,g,k), and carpels (d,h,l) of WT (b-d), *fa^pi^* (f-h), and *fa^bif^*(j-l) flowers. Number of organs per whorl in *fa^pi^* (m) and *fa^bif^* (n) flowers. ns, not significant difference; * significant differences at *P* < 0.05. (Student’s t-test).

**Fig. 4.**
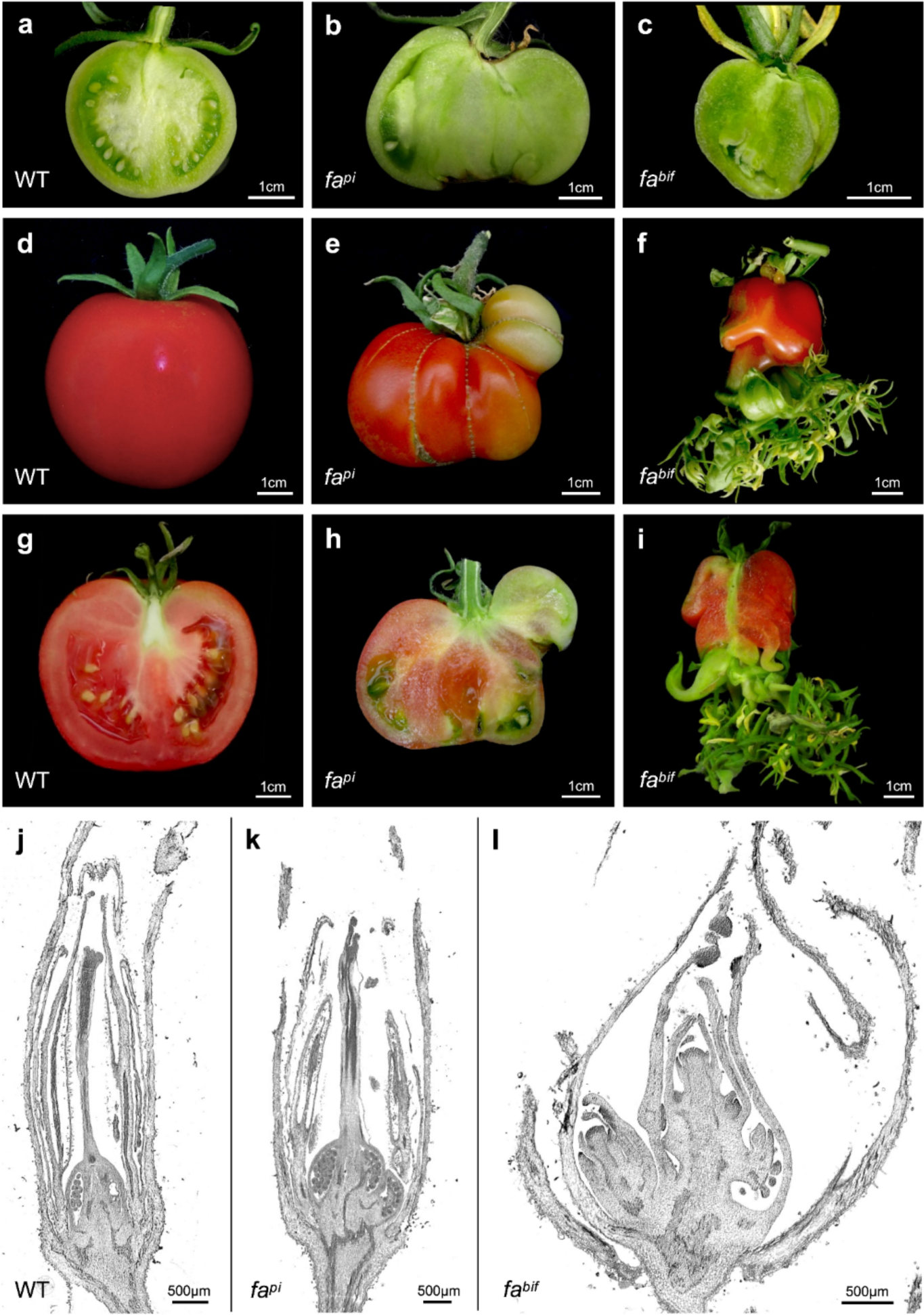
Fruit development in the *fa^bif^* hypomorphic allele reveals that *FA* plays a role in floral determinacy. Representative longitudinal open fruits of wild-type (WT) (a), *fa^pi^*(b), and *fa^bif^* (c) plants at the mature green stage. Closed fruits (d-f) and longitudinal open fruits (g-i) of WT (d,g), *fa^pi^*(e,h), and *fa^bif^* (f,i) plants at the red ripe stage. Histological sections of flowers at anthesis stage of WT (j), *fa^pi^* (k), and *fa^bif^* (l) plants.

A detailed comparative analysis of *fa*, *fa^pi^*, and *fa^bif^* was performed to evaluate the impact of allelic variation at the *FA* locus on the reproductive development of tomato. Regarding flowering time, assessed as the number of leaves before the first inflorescence, a significant delay in flowering transition was observed in *fa* (7.63 ± 1.55), *fa^pi^* (7.06 ± 1.03), and *fa^bif^*(5.34 ± 0.73) mutants compared with their respective wild-type (WT) genetic backgrounds (5.67 ± 1.23, 5.47 ± 0.72, and 4.92 ± 0.81, respectively) (Fig. 1m). In addition, the early morphogenesis of WT and mutant inflorescences was compared in-depth through microscopic observations. At the time of floral transition, the shoot apex of WT plants developed into the primary IM, which subsequently gave rise to the first FM, while a lateral IM formed adjacent to it. As the FM matured into a flower, the secondary IM produced a new lateral IM before transitioning to the next FM (Fig. 1a,b). This growth sequence continued repeatedly, resulting in the formation of an inflorescence with a zigzag pattern (Fig. 1c). In the *fa* null mutant (amorph allele), the shoot apex gave rise to the development of a primary IM that remained indeterminate and repeatedly initiated secondary IMs (inflorescence shoots). However, these IMs were unable to transition to the FM fate and instead reverted to vegetative growth (Fig. 1d,e). This developmental pattern eventually led to the formation of a complex inflorescence with a leafy appearance, lacking any floral structures (Fig. 1f,n). In contrast, *fa^pi^* and *fa^bif^*hypomorphic mutants developed branched inflorescences with both leafy and floral organs (Fig. 1i,l). In the *fa^pi^* mutant, the shoot apex gave rise to the inflorescence by developing a primary IM, which formed additional IMs before one of them transitioned into the first FM (Fig. 1g,h). A similar developmental pattern was observed in the *fa^bif^* mutant, although in this case, the IM-to-FM transition was even more delayed than in the *fa^pi^* mutant, resulting in the development of a large number of inflorescence shoots before the formation of the first FM (Fig. 1j,k). These sequential delays in the IM-to-FM transition eventually led to the formation of highly branched inflorescence with an increased number of flowers in both *fa^pi^* and *fa^bif^*mutants (Fig. 1i,l,n), although most of these flowers showed organ abnormalities, as described in the following section.

To further assess the role of *FA* in flowering time and inflorescence branching, we developed *FA* hypermorph alleles by constitutive overexpression of the *FA* gene. Three independent second-generation transgenic (TG2) progeny expressing a *35S:FA* transgene were generated. These plants flowered very early, just after the formation of 1.67 ± 0.93 leaves (Fig. 2a-d), whereas floral transition of WT plants occurred after the formation of 7.88 ± 1.26 leaves. Furthermore, the overexpression of *FA* resulted in a reduction in the number of flowers per inflorescence. Concretely, *35S:FA* plants formed 2.78 ± 1.36 flowers per inflorescence, whereas WT plants developed 8.93 ± 2.66 flowers (Fig. 2e). In addition, we used RNA interference (RNAi) approach to generate knockdown alleles of *FA*. Three first-generation *RNAi:FA* independent transgenic (TG1) lines were obtained, but none of these lines produced fruits, making it impossible to evaluate their TG2 progeny. Therefore, five clonal replicates of each TG1 were generated to evaluate their *FA* expression levels and phenotypes. *FA* transcripts were found to be downregulated in the reproductive transition meristems of the RNAi plants, although the RNAi lines exhibited varying degrees of *FA* repression. Specifically, the *RNAi:FA-1* and *RNAi:FA-3* lines showed a significant reduction in *FA* expression compared to the WT, while intermediate expression levels were observed in plants from the *RNAi:FA-2* line (Fig. 2i). According to the expression results, the inflorescences of the *RNAi:FA-1* and *RNAi:FA-3* lines presented more pronounced alterations in their development compared to those observed in *RNAi:FA-2* inflorescences. The phenotype of *RNAi:FA-1* and *RNAi:FA-3* inflorescences resembled that of the *fa* amorph allele, characterized by the formation of indeterminate leafy inflorescences with IMs that are unable to transition to the FM fate and eventually reverted to vegetative growth. Conversely, a phenotype similar to the *fa^pi^* hypomorphic mutant was observed in *RNAi:FA-2* plants, which displayed branched inflorescences bearing both leaves and flowers with aberrant floral organs (Fig. 2f-h, j). Since these TG1 *RNAi:FA* plants were derived from regenerated transformed cells, evaluation of flowering time was precluded, limiting the phenotypic assessment to inflorescence development. Taken together, these results indicate that *FA* not only participates in flowering transition but also plays a role in IM maturation and termination, thereby regulating the inflorescence architecture.

### Hypomorphic alleles of *FA* develop flowers with homeotic conversions and reduced organ number

Unlike the *fa* null mutant, whose IMs were unable to give rise FMs and reverted to a vegetative developmental program, the *fa^pi^* and *fa^bif^* hypomorphic mutations showed weaker effects and formed FMs that matured into flowers. However, different degrees of homeotic transformations were observed in their floral organs. In *fa^pi^* flowers, sepals were longer than in WT, while the petals were shorter and narrower (Fig. 3a, e). Additionally, *fa^pi^* flowers usually exhibited leaf-like sepals on the flower pedicels along with the sepals that developed concentrically in the first whorl. Compared to WT, *fa^pi^* petals showed distorted shape, although no homeotic conversions were found (Fig. 3b, f). In contrast, WT stamen morphology (Fig. 3c) was almost never observed in *fa^pi^* flowers. The *fa^pi^* stamens were transformed into chimeric organs with anther tissues mixed with cells bearing petaloid identity in the middle sections. Carpelloid tissues were also found in the basal regions, where ectopic ovules were observed (Fig. 3g). Furthermore, whereas the stigma, style and carpel tissues were clearly observed in WT pistils (Fig. 3d), *fa^pi^*pistils usually showed varying degrees of homeotic changes; in most cases, they were partially transformed into petals, giving them a yellow-green color (Fig. 3e, h). Stigma and style tissues were rarely found in *fa^pi^* pistils, although papillary cells were occasionally developed in the distal part (Fig. 3e).

Regarding *fa^bif^* flowers, also had longer sepals and petals that were generally smaller in width and length than those of WT. In addition, leaf-like sepals were commonly found on the flower pedicels (Fig. 3i). Although *fa^bif^* petals were deformed, no homeotic transformations were detected (Fig. 3j). However, stamens and pistils displayed transformations in organ identity similar to those observed in the *fa^pi^* mutant. The *fa^bif^* stamens were chimeric organs with sections exhibiting petal morphology, as well as growing structures with cellular identity similar to that observed in the WT styles (Fig. 3d, k). Likewise, ectopic ovules were developed in the *fa^bif^* stamens (Fig. 3k). Furthermore, *fa^bif^* flowers developed pistils that also displayed partial homeotic transformations (Fig. 3l), although these were less severe than those observed in *fa^pi^* pistils. Frequently, *fa^bif^* pistils exhibited lateral sections with petaloid identity and a yellow-green color (Fig. 3i, l).

Additionally, the number of floral organs was assessed in *fa^pi^*and *fa^bif^* mutants (Fig. 3m, N). The number of sepals was significantly lower and higher than WT in *fa^pi^*and *fa^bif^* flowers, respectively. In *fa^pi^* flowers, the number of true sepals was higher compared to *fa^bif^*. In contrast, *fa^bif^* developed a greater number of leaf-like sepals than *fa^pi^*. No differences were found between WT and mutant flowers in the number of petals and carpels. Furthermore, both *fa^pi^* and *fa^bif^* flowers developed a significantly reduced number of stamens (Fig. 3m, n). Taken together, these results demonstrate that *FA* is required for the establishment of floral organ identity and contributes to the specification of the number of organs in each whorl, with a primary impact on the stamens.

### *fa^bif^* alters floral determinacy giving rise to indeterminate fruits

Despite the abnormalities found in the floral development of *fa^pi^*and *fa^bif^* mutants, both were able to produce fruits. In WT plants, seeds are fully developed and embedded in the locular gel at the mature green fruit stage (Fig. 4a). In contrast, the locular cavities in the *fa^pi^* and *fa^bif^* fruits were predominantly filled with placental tissue. Remarkably, while seeds occasionally developed in *fa^pi^*fruits, this feature was never observed in *fa^bif^* (Fig. 4b, c). The WT plants developed determinate fruits with a round shape and a uniform red color at the red ripe stage (Fig. 4d, g). At this developmental stage, the *fa^pi^*mutant also produced determinate fruits, although they displayed a distorted shape and uneven ripening (Fig. 4e, h). In contrast, *fa^bif^*fruits were indeterminate, with broken pericarp, and ectopic shoots emerged from these fruits (Fig. 4f, I). These ectopic shoots led to the formation of leaf-like structures and flowers with a phenotype similar to that observed in the flowers previously developed by *fa^bif^*. Histological sections of flowers at the anthesis stage were performed to further characterize fruit ontogeny. Unlike the WT and *fa^pi^* flowers, in which ovules were developed within carpels (Fig. 4j, k), *fa^bif^* flowers generated pistil walls, but inside them, instead of ovules, new vegetative sympodial meristems were formed (Fig. 4l). The development of indeterminate flowers in *fa^bif^*reveals that *FA* has a role in specifying floral determinacy, enabling the cessation of stem cell activity in the FM to allow the proper formation of flowers and fruits.

### CRISPR/Cas9 genome editing to expand *FA* allelic series

To further explore the functional role of *FA* in tomato reproductive development, genome editing approach was used to generate new alleles through disrupting the coding and regulatory regions of the *FA* gene. Mutations in the *FA* coding sequence (*fa^CR-cds^*) were engineered using the CRISPR/Cas9 system with a single guide RNA (sgRNA) targeting its first exon (Fig. 5a). Three independent TG1 diploid lines were identified, each carrying edited alleles in the region targeted by the sgRNA in the homozygous state. The *fa^CR-cds-1^* plant carried a single A deletion, whereas four-nucleotide deletion was found (-CCAA and -AAGC, respectively) in *fa^CR-cds-2^* and *fa^CR-cds-3^* plants (Fig. 5a). The phenotypic characterization of these three TG1 lines was limited to the level of inflorescence development, since they were obtained through *in vitro* regeneration of transformed cells, which precludes the evaluation of flowering time. In all cases, the shoot apex of the *fa^CR-cds^* lines generated an indeterminate IM that continuously produced secondary IMs. However, these IMs did not progress to FM fate and instead reverted to vegetative growth. Thus, *fa^CR-cds^* displayed a phenotype almost identical to that of the *fa* amorph allele, with leafy inflorescences unable to develop flowers (Fig. 5b,c).

**Fig. 5.**
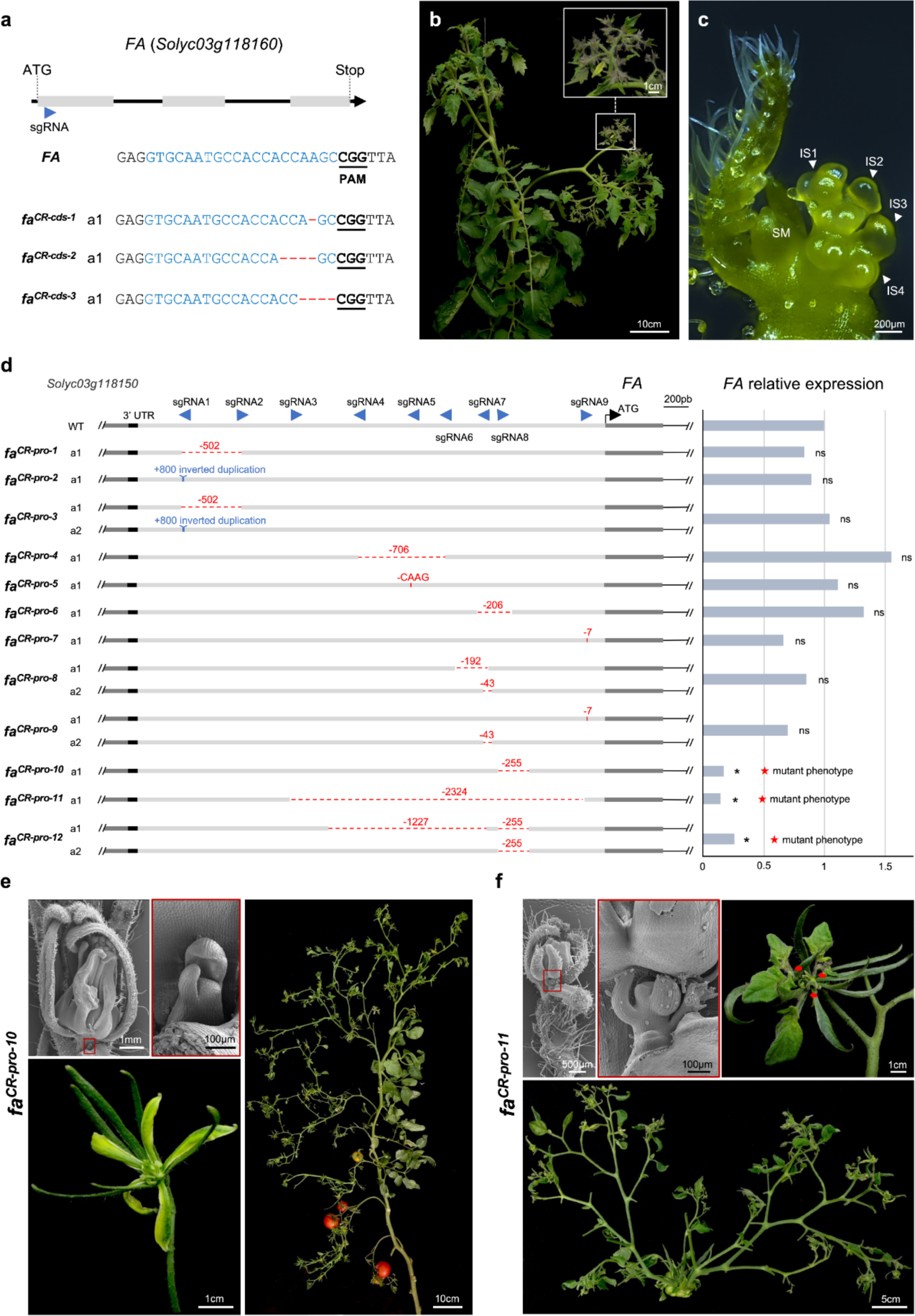
Characterization of CRISPR/Cas9 lines targeting the coding (*fa^CR-cds^*) and regulatory (*fa^CRpros^*) regions of the *FA* gene. (a) Scheme illustrating single guide RNA (sgRNA) targeting the *FA* coding sequence (blue arrowheads). *fa^CR-cds^* alleles identified by cloning and sequencing PCR products from the *FA*-targeted region. Red dashed lines indicate InDel mutations, blue font highlights the sgRNA target, black bold and underlined letters indicate the protospacer adjacent motif (PAM) sequence. a, allele. (b) Representative *fa^CR-cds^* inflorescence architecture. (c) Stereoscopic images illustrating inflorescence development of *fa^CR-cds^* plants. SM, sympodial meristem; IS, inflorescence shoot. (d) Schematic representation of mutations identified in the *FA* promoter region in *fa^CR-pro^* lines. Deletions (–) and insertions (+) indicated by numbers or letters. qRT-PCR analysis of *FA* expression from WT and *fa^CR-pro^* reproductive meristems. ns, not significant difference; * significant differences at *P* < 0.05. (Student’s t-test). Representative phenotypic alterations identified in *fa^CR-pro-10^* (E) and *fa^CR-pro-11^* (F) lines.

In addition, three CRISPR/Cas9 vectors, each containing three sgRNAs, were constructed to generate cis-regulatory mutations by targeting the 4-kb promoter region immediately upstream of the *FA* coding sequence (*fa^CR-pro^*). We identified 12 *fa^CR-pro^* lines carrying homozygous or biallelic edited alleles in the *FA* promoter region. For example, the *fa^CR-pro-1^*and *fa^CR-pro-2^* lines were homozygous for a 502-bp deletion and an 800-bp inverted duplication, respectively. In contrast, the *fa^CR-pro-3^* line was biallelic for the two alleles mentioned above (Fig. 5d). In total, eleven different *fa^CR-pro^* alleles were identified, representing a variety of mutation types, including large deletions, inversions, and small indels (Fig. 5d). We also evaluated the expression of *FA* in reproductive transition meristems of the *fa^CR-pro^* lines. Among these lines, *fa^CR-pro-10^*, *fa^CR-pro-11^*, and *fa^CR-pro-12^*showed a significant reduction in *FA* expression levels compared to the WT (Fig. 5d). The *fa^CR-pro-10^* and *fa^CR-pro-11^* lines were homozygous for a 255-bp deletion and a 2324-bp deletion, respectively, while the *fa^CR-pro-12^* line was biallelic, carrying the 255-bp deletion on both alleles and, an additional 1227-bp deletion on one of them, (Fig. 5d). Remarkably, these three lines displayed a phenotype similar to that of the *fa^pi^* and *fa^bif^* hypomorphic alleles. The *fa^CR-pro-10^* and *fa^CR-pro-12^* lines displayed an identical phenotype characterized by compound inflorescences with leaves and flowers with diverse degrees of homeotic conversions between floral organs, primarily affecting the third and fourth whorls (Fig. 5e). A distinctive feature of *fa^CR-pro-10^* and *fa^CR-pro-12^* flowers was the development of vegetative sympodial shoots at the sepal axil, although these shoots did not continue to grow on mature flowers (Fig. 5e). In the *fa^CR-pro-11^*branched inflorescences, flowers frequently developed leaf-like organs, and vegetative sympodial shoots also emerged at the sepal axil. However, unlike *fa^CR-pro-10^* and *fa^CR-pro-12^*, these vegetative sympodial shoots progressed in their development, leading to the formation of new inflorescences at the axil of the fruit sepals (Fig. 5f). These results support that *FA* is needed to specify inflorescence architecture and floral organ identity, but it is also involved in floral meristem determinacy, as a reduction in its expression levels prevents the cessation of meristematic activity in flowers.

### Molecular characterization and sequence analysis of the *FA* allelic series

The comparison of the orthologous LFY sequences from multiple species revealed that the amino acid residues of the SAM and DBD domains are highly conserved throughout LFY evolution (Supplementary Fig. S1). The *fa* amorph allele is due to a 16-bp deletion resulting in a frameshift mutation that leads to a truncated FA protein lacking the DBD domain (Molinero-Rosales et al. 1999; Fig. 6). To determine the molecular nature of the *fa^pi^* and *fa^bif^*hypomorphic mutations, the *FA* genomic region was sequenced. In *fa^pi^*, a T/C transition was found at the 302nd nucleotide position of the *FA* coding sequence, generating a non-synonymous mutation from Met to Thr at position 101 (M101T). In the *fa^bif^* mutant, a G/C transversion was found 249-bp downstream of the *FA* translation start codon, giving rise to another non-synonymous mutation from Gly to Arg at position 84 (G84R). Both *fa^pi^* and *fa^bif^* mutations were located at the SAM oligomerization N-terminal domain, affecting highly conserved positions (Fig. 6, Supplementary Fig. S1). Concerning CRISPR/Cas9 lines targeting the *FA* coding sequence, the single A deletion identified in the *fa^CR-cds-1^* line gives rise to a frameshift mutation at the 23rd position of the FA protein sequence. The four-nucleotide deletions found in *fa^CR-cds-2^* and *fa^CR-cds-3^* lines (-CCAA and -AAGC, respectively) also result in frameshift mutations, both affecting the 22nd amino acid position. Unlike *fa* amorph allele, the FA truncated proteins from the *fa^CR-cds^* alleles lack both the SAM and DBD domains (Fig. 6).

**Fig. 6.**
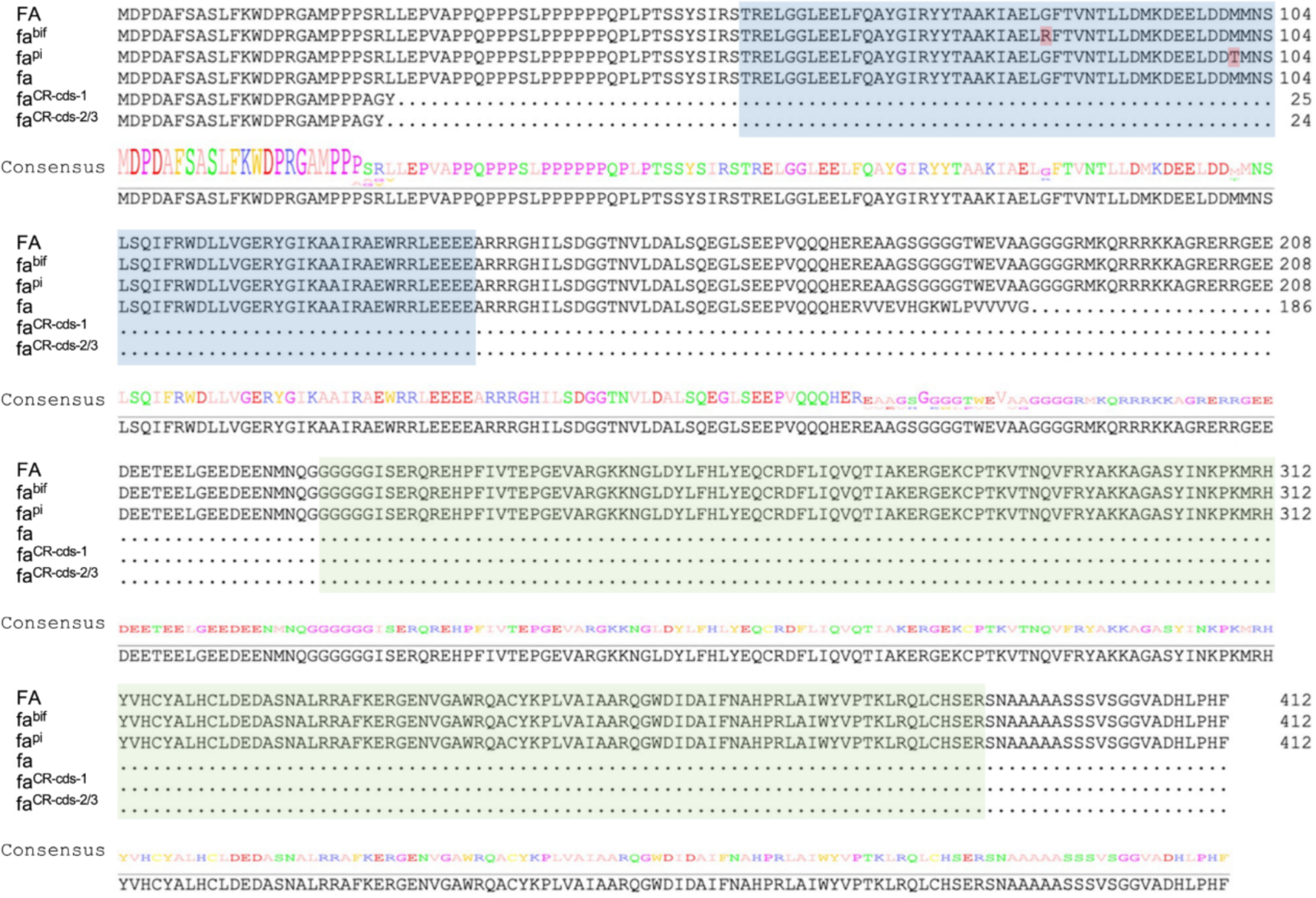
Amino acid sequence alignment of the wild-type (WT) FA protein with the predicted proteins from the *fa* mutant alleles. The sequence alignment and consensus sequence logo (at the bottom) were generated by Jalview. The consensus sequence is displayed in logo format with the amino acid expectation illustrated by relative letter size at each position. The SAM and DBD domains are indicated in blue and green colors, respectively. The residue positions changed by *fa^pi^* and *fa^bif^*mutations are shown in red.

### *SlWUS* and *TAG1* expression during flower development

Floral determinacy involves the cessation of stem cell activity through the repression of the meristem identity gene *SlWUS* (Bollier et al. 2018; Castañeda et al. 2022). Therefore, we further investigated the function of *FA* in FM determination by studying the temporal expression of *SlWUS* during flower development. qRT-PCR analysis of *SlWUS* expression was conducted in hypomorphic mutants, capable of developing flowers, and their respective WT genetic backgrounds at seven developmental stages, ranging from reproductive meristem (RM) to anthesis + 2 days (AD+2; see Material and Methods). In the *fa^pi^* mutant, which showed no visible alterations in FM determinacy, *SlWUS* expression was significantly downregulated in floral buds 1 (FB1), whereas its transcript levels were upregulated in AD and AD+2 (Fig. 7a). The most notable differences in the *SlWUS* expression pattern were found in *fa^bif^*mutant, which showed clear defects in FM termination during fruit development. *SlWUS* transcripts decreased in *fa^bif^* from RM to pre-anthesis (PA) stages. In contrast, while *SlWUS* expression levels were almost undetectable in WT flowers at A and A+2 stages, they were significantly increased in *fa^bif^* flowers (Fig. 7c). A similar *SlWUS* expression pattern was observed in the *fa^CR-pro-10^* edited line. The downregulation of *SlWUS* was observed from RM to FB2 stages. However, no significant differences were observed at PA and A stages, although *SlWUS* transcript levels were significantly higher at AD+2 stage (Fig. 7e).

**Fig. 7.**
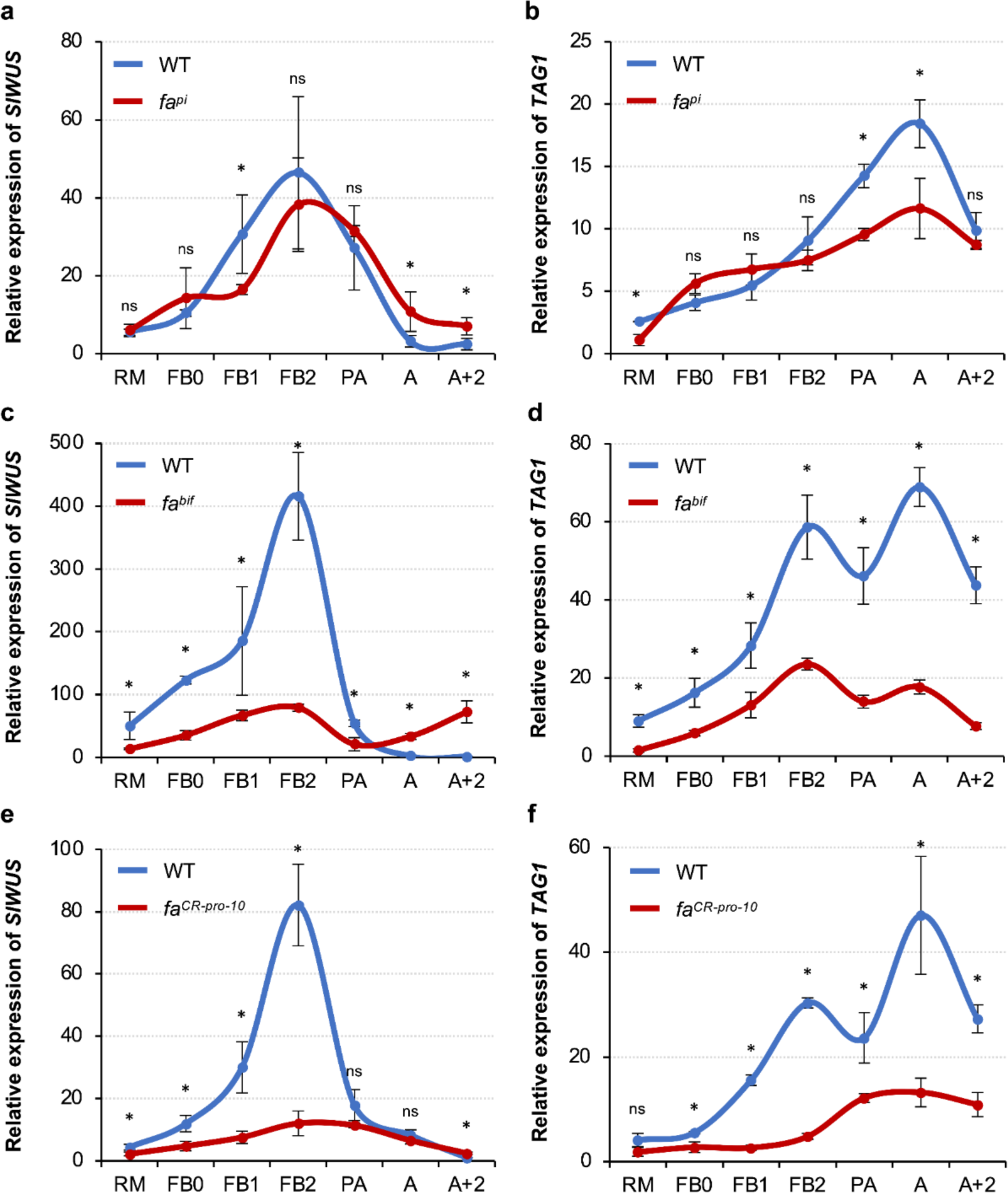
Expression pattern of *SlWUS* and *TAG1* during flower development. Relative *SlWUS* expression in *fa^pi^* (a), *fa^bif^* (c) and *fa^CR-pro-10^* (e) plants compared to their respective wild-type (WT) genetic backgrounds. *TAG1* relative expression in *fa^pi^* (b), *fa^bif^* (d) and *fa^CR-pro-10^*(f) plants. RM, reproductive meristem; FB0, floral buds of 1.0–2.9 mm in length; FB1, floral buds of 3.0–4.9 mm in length; FB2, floral buds of 5.0–7.9 mm in length; PA, flowers at the pre-anthesis stage; A, flowers at the anthesis stage; AD+2, flowers at the anthesis stage + 2 days; ns, not significant difference; * significant differences at *P* < 0.05. (Student’s t-test).

Molecular mechanism mediated by WUS expression that promotes flower development has been further studied in Arabidopsis. In this model species, LFY cooperates with WUS to induce *AG* transcription in flower buds at stage 3 (Lenhard et al. 2001; Lohmann et al. 2001). Nevertheless, no information regarding this mechanism has been reported in tomato. Therefore, we also evaluated the expression of *TAG1*, the tomato AG homologue (Pnueli et al. 1994; Gimenez et al. 2016), during the reproductive stages of tomato plants. The expression of *TAG1* in *fa^pi^* mutant was altered in the RM, PA, and A stages, showing lower transcript levels than in the WT genetic background (Fig. 7b). In *fa^bif^* mutant, *TAG1* transcripts were downregulated in all assessed developmental stages, from RM to A+2 (Fig. 7d). A similar expression pattern was found in the *fa^CR-pro-10^* edited line. Except for the RM stage, where no significant differences with respect to WT were observed, *TAG1* expression was downregulated from FB to A+2 (Fig. 7e). Taken together, these results indicate that FA is directly or indirectly involved in the regulation of *SlWUS* and *TAG1* expression, and that the observed abnormalities in the temporal expression patterns of these two genes may be responsible for the floral indeterminacy phenotypes observed in mutant plants.

## Discussion

Significant advancements have been achieved in unraveling the intricate mechanisms that orchestrate floral transition and floral growth and differentiation, much of which stems from studies conducted in the model species *A. thaliana*, where LFY acts as a pioneer transcription factor. LFY plays a pivotal role in the reproductive phase by modulating chromatin structure, which renders DNA accessible to other transcription factors and regulatory proteins, thereby influencing gene expression patterns and ultimately determining floral fate (Jin et al. 2021; Lai et al. 2021; Yamaguchi 2021). Structure-function studies have revealed that LFY interacts with semi-palindromic 19-bp *cis*-elements through its highly conserved DBD domain situated at the C-terminal region. This domain adopts a helix-turn-helix fold, enabling dimerization on DNA (Moyroud et al. 2011; Winter et al. 2011; Sayou et al. 2016). Furthermore, although the SAM oligomerization N-terminal domain does not make directly contact with DNA, it provides LFY with the capability to access closed chromatin regions, thereby modulating the DNA binding landscape of LFY (Sayou et al. 2016). With respect to the tomato LFY orthologue, although structure-function studies on FA have not been conducted, the comparative analysis performed herein on the *FA* allelic series may provide insights into the functional significance of the SAM and DBD domains in the reproductive phase of tomato. The absence of the DBD domain in the *fa* and *fa^CR-cds^* amorph alleles (Fig. 6) results in leafy inflorescences incapable to develop flowers (Fig.1 d-e; Fig. 5b,c). Conversely, the *fa^pi^* and *fa^bif^* hypomorphic alleles, which carry non-synonymous mutations at the SAM oligomerization N-terminal domain (Fig. 6), produce flowers with varying degrees of homeotic transformation (Fig. 3e-l). Unlike the *fa^CR-cds^* alleles, the FA protein produced by the *fa* allele retains the SAM domain intact (Fig. 6). However, despite this difference, these genotypes exhibited almost identical phenotypes. These findings suggest that while the DBD domain may play a crucial role in determining FM identity, the SAM domain may be involved in both specifying and maintaining floral organ identity. In accordance with this proposal, the *fa^lfi^* mutant, which carries a 12-bp deletion resulting in an in-frame mutation located within the DBD domain, is unable to produce flowers and instead develops inflorescences with fleshy carpeloid leaves (Kato et al. 2005). To our knowledge, the involvement of the SAM and DBD domains in specific processes of Arabidopsis reproductive development has not been addressed. In the *lfy-1* mutant (Q32stop), both the SAM and DBD domains are absent, similar to the *fa^CR-cds^* alleles. Conversely, the *lfy-7* mutant (Q187stop) resembles the *fa* allele, with only the DBD domain missing in their respective proteins (Weigel et al. 1992). Similar to observations in tomato, both *lfy-1* and *lfy-7* alleles display strong mutant phenotype, characterized by the failure of IMs to transition into FMs. Other mutants carrying non-synonymous mutations in the DBD domain have been described, such as the *lfy-3* (T244M), *lfy-4* (E238K), *lfy-5* (P240L), *lfy-9* (R331K), *lfy-20* (N306D) and *lfy-28* (P308L) alleles, but all of them show weak or intermediate mutant phenotypes (Weigel et al. 1992; Hamès et al. 2008). Although additional studies are required, the overall results indicate the essential role of the DBD domain in determining FM identity in both Arabidopsis and tomato.

The evaluation of the *FA* allelic series underscores its multifaceted role in tomato reproductive development, exerting control over both early and late stages of floral ontogeny. The involvement of *FA* in promoting floral transition, FM identity, and floral organ identity has already been stated above (Allen and Sussex 1996; Molinero-Rosales et al. 1999; Kato et al. 2005; Olimpieri and Mazzucato 2008). However, the extreme compound inflorescence phenotype caused by the loss of functionality or downregulation of the *FA* gene, along with the fact that the overexpression of *FA* accelerates flowering and converts a multi-flowered inflorescence into a single flower, suggest an additional function for *FA* in IM maturation and termination. Considering these findings, it is plausible to hypothesize that varying levels of *FA* expression could modulate the branching and the number of flowers formed per inflorescence, which could be of potential interest from an agronomical point of view. To further validate this hypothesis, we attempt to modify *FA* expression levels by editing its *cis*-regulatory elements using CRISPR/Cas9 technology, a strategy that has been demonstrated to be effective in tomato (Rodríguez-Leal et al. 2017) and maize (Liu et al. 2021) to generate quantitative variation in yield-related traits. However, we observed only two distinct *FA* expression patterns in *fa^CR-pro^* edited lines. One group of lines exhibited no significant differences in expression compared to the WT and displayed no phenotypic alterations, while the other group displayed markedly reduced *FA* expression levels, resulting in an inability to develop normal flowers (Fig. 5d-f). Although we were unable to find subtle quantitative changes in inflorescence branching, the molecular characterization of the *fa^CR-pro^* lines has unveiled insights into a specific promoter region that appears to play an important role in promoting *FA* expression. Remarkably, the *fa^CR-pro-10^*, *fa^CR-pro-11^*, and *fa^CR-pro-12^*lines share a common 255-bp deletion that results in a drastic decrease in *FA* transcript levels (Fig. 5d). Hence, this region could serve as a potential target for regulators of *FA*, whose study would be of great interest in furthering our understanding of tomato reproductive development.

The role of *FA* in specifying floral determinacy is evidenced by the phenotypes of *fa^bif^* (Fig. 4l) and *fa^CR-pro^*(Fig. 5e, f) plants, which develop new sympodial meristems within their flowers, revealing their inability to arrest stem cell activity in the FM. In Arabidopsis, LFY is indirectly involved in the repression of the stem cell identity gene *WUS* through the activation of the C-class MADS-box gene *AG* (Lenhard et al. 2001; Lohmann et al. 2001). This activation leads to the cessation of the stem cell maintenance program via distinct pathways: directly, through chromatin remodeling and the recruitment of the Polycomb group (PcG) protein TERMINAL FLOWER2/LIKE HETEROCHROMATIN PROTEIN1 (TFL2/LHP1) at the *WUS* locus (Guo et al. 2018), and indirectly, through the transcriptional induction of two key targets, *KNUCKLES* (*KNU*) and *CRABS CLAW* (*CRC*). Both *KNU* and *CRC* act through independent pathways to synergistically regulate *WUS* repression, and thereby trigger FM termination (Sun et al. 2009, 2014, 2019; Yamaguchi et al. 2017, 2018). Previously, we have shown, that tomato SlCRC paralogues interact with members of the chromatin remodeling complex responsible for the epigenetic regulation of *SlWUS* expression (Castañeda et al. 2022). In this epigenetic silencing mechanism, INHIBITOR OF MERISTEM ACTIVITY (SlIMA) and SlKNU facilitate the recruitment of TOPLESS1 (SlTPL1) and HISTONE DEACETYLASE1 (SlHDA1) to form a transcriptional repressor complex at the *SlWUS* locus (Bollier et al. 2018). The role of FA in repressing *SlWUS* to terminate stem cell activity has remained unexplored until now. Our expression analysis in hypomorphic mutants revealed that the smallest differences in *SlWUS* expression levels were found in the *fa^pi^* mutant (Fig. 7a), where no alterations in FM determinacy were observed. In contrast, *SlWUS* transcripts were less abundant in *fa^bif^* and *fa^CR-pro-10^* plants compared to WT plants during floral development, while its expression increased at the A+2 stage (Fig. 7c,e). This increase could possibly explain the emergence of new sympodial meristems, since the stem cell identity gene *SlWUS* remained active. Similarly, *TAG1* expression was less affected in *fa^pi^* flowers compared to *fa^bif^* and *fa^CR-pro-10^* flowers. Furthermore, unlike *SlWUS*, the transcript abundance of *TAG1* was lower in *fa^bif^* and *fa^CR-pro-10^* flowers across all floral stages evaluated (Fig. 7b,d,f). These findings are consistent with those reported by Kato et al. (2005), who observed that *TAG1* expression was inhibited in *fa^lfi^* inflorescences. Collectively, these data suggest that FA may directly trigger the expression of *TAG1*, as LFY does with *AG* in Arabidopsis (Lenhard et al. 2001; Lohmann et al. 2001). However, FA may not act alone, since *TAG1* is still expressed in the absence of *FA* function. Thus, the subdued expression levels of *TAG1* could contribute to the deregulation of *SlWUS* during floral development, which, in turn, would promote the appearance of the fruit-into-fruit phenotypes characteristic of *fa^bif^* and *fa^CR-pro-10^*.

Taken together, our findings reveal the multidimensional function of *FA* in regulating the reproductive phase of tomato. In addition to promoting floral transition and specifying the identity of FMs and floral organs, *FA* also exerts regulatory control over the expression of *TAG1* and *SlWUS*, underscoring its essential role in promoting carpel development and suppressing floral stem cell activity. Thus, our study highlights the potential of using allelic series of mutants as powerful tools for unraveling gene functions and deciphering the intricate molecular basis of biological processes.

## Data availability

The authors affirm that all data necessary for confirming the conclusions of the article are present within the article, figures, and tables.

## Acknowledgments

The authors would like to thank Campus de Excelencia Internacional Agroalimentario (CeiA3) for providing research facilities.

## Supplementary Material

**Supplementary Fig. S1.**
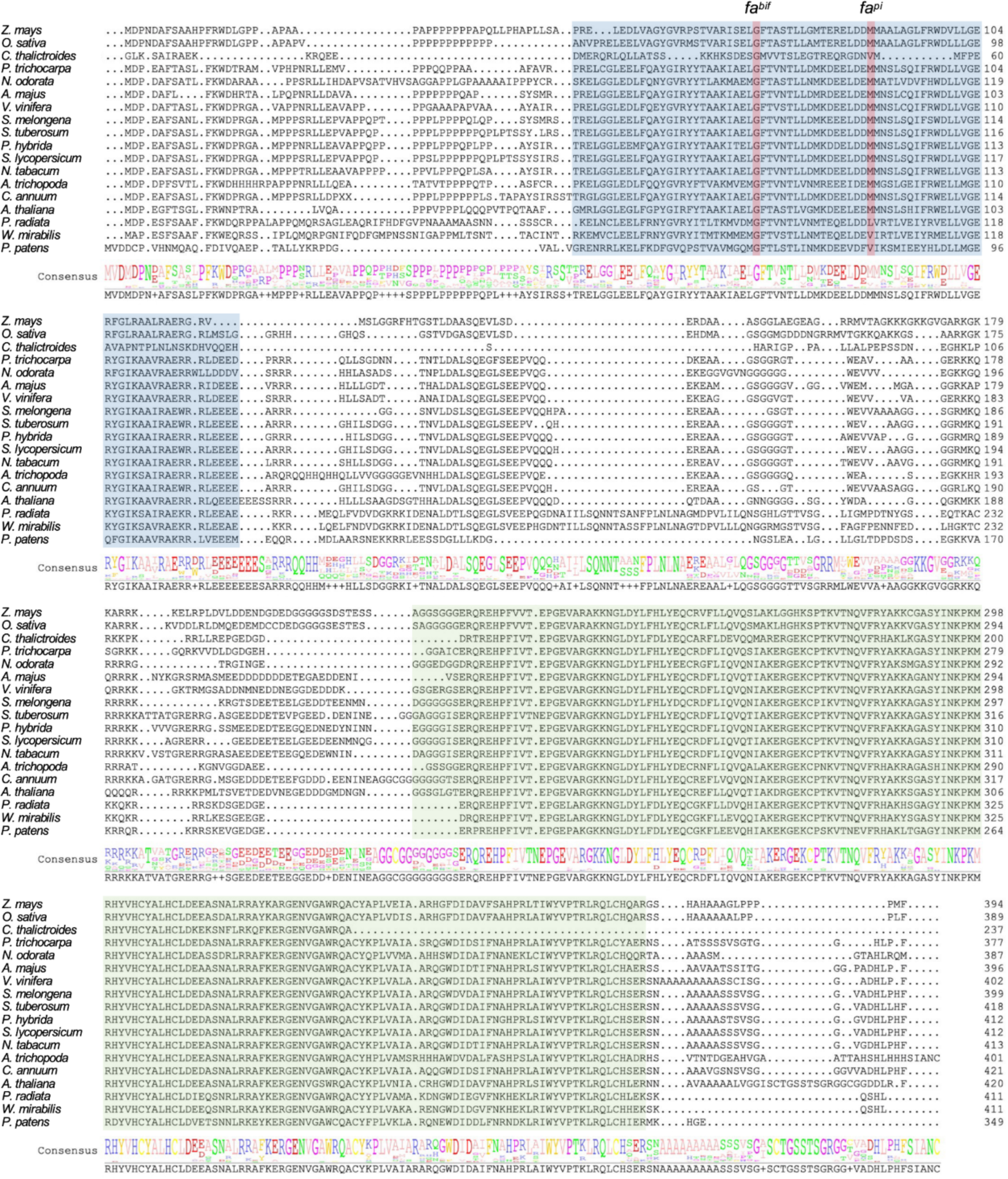
Amino acid sequence alignment of LEAFY orthologues from multiple species. Jalview (https://www.jalview.org/) was used to generate both the sequence alignment and the consensus sequence logo (located at the bottom). The consensus sequence is presented in a logo format, where the amino acid expectation is depicted by the relative letter size at each position. The SAM and DBD domains are highlighted in blue and green colors, respectively. The residue positions affected by *fa^pi^* and *fa^bif^* non-synonymous mutations are shown in red.

**Supplementary Table S1.**
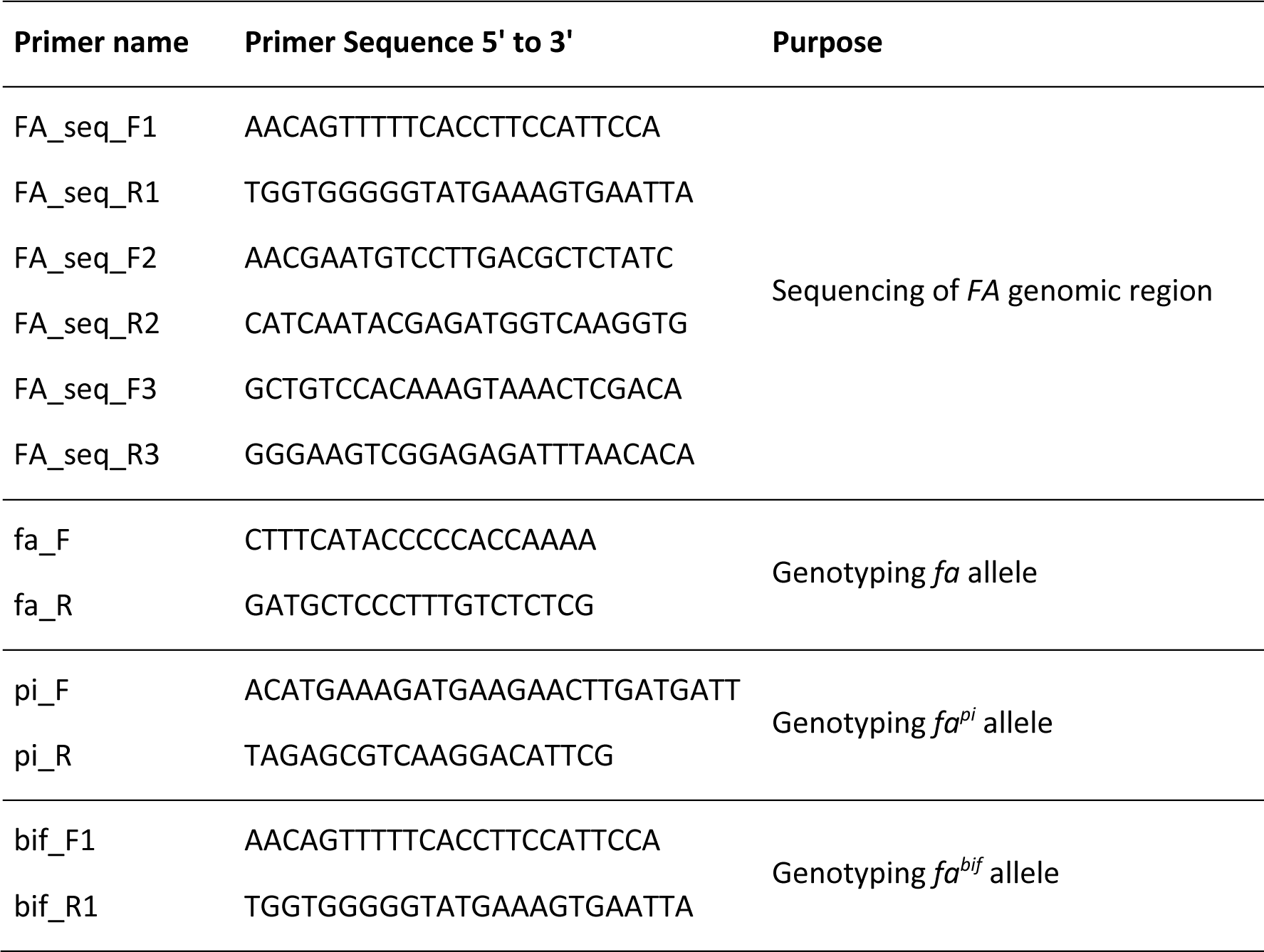
Primer sequences used to sequence the *FA* genomic region and genotype mutant alleles.

**Supplementary Table S2.**
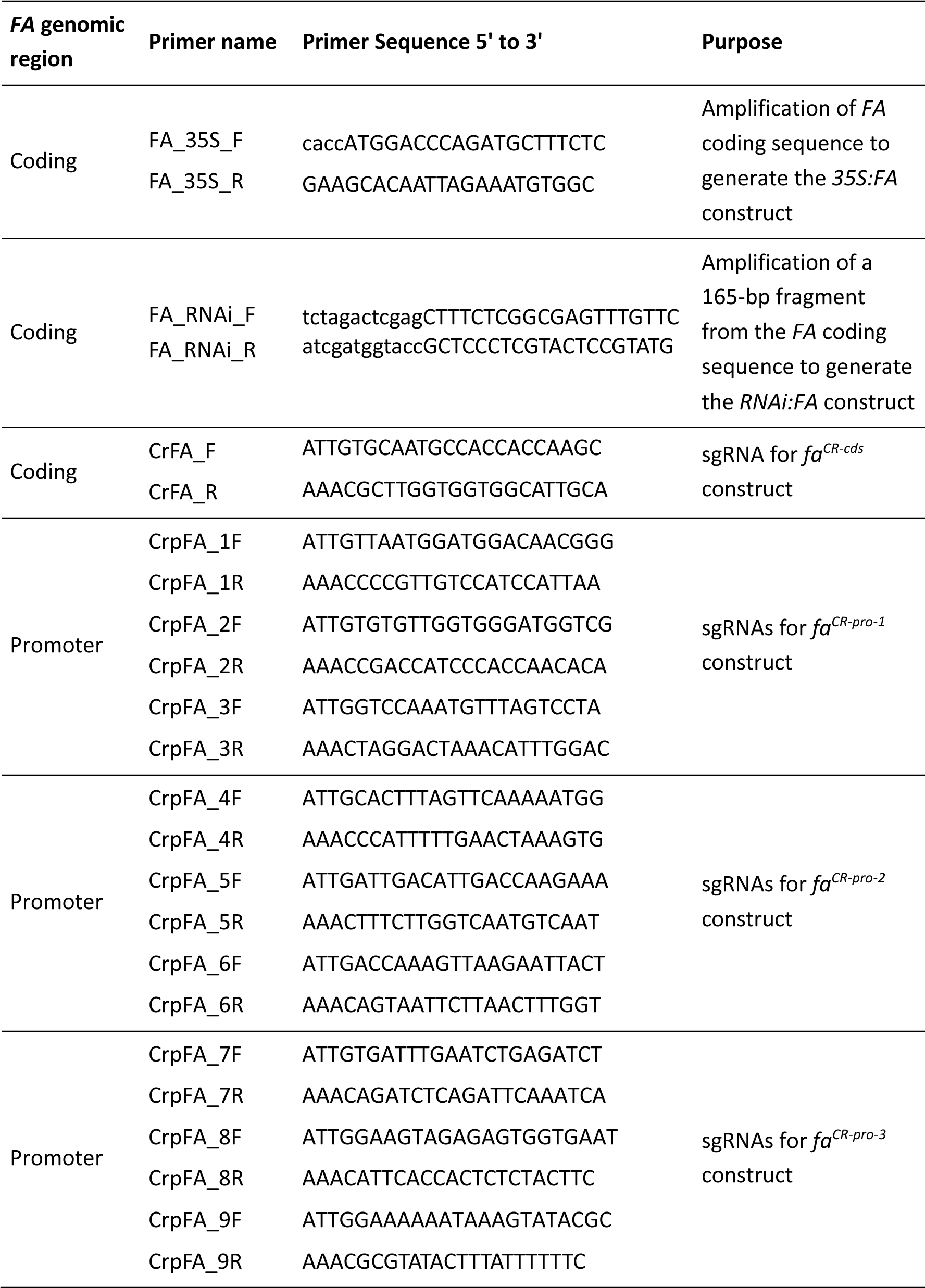
Primer sequences used to generate overexpression, RNAi and CRISPR/Cas9 vectors.

**Supplementary Table S3.**
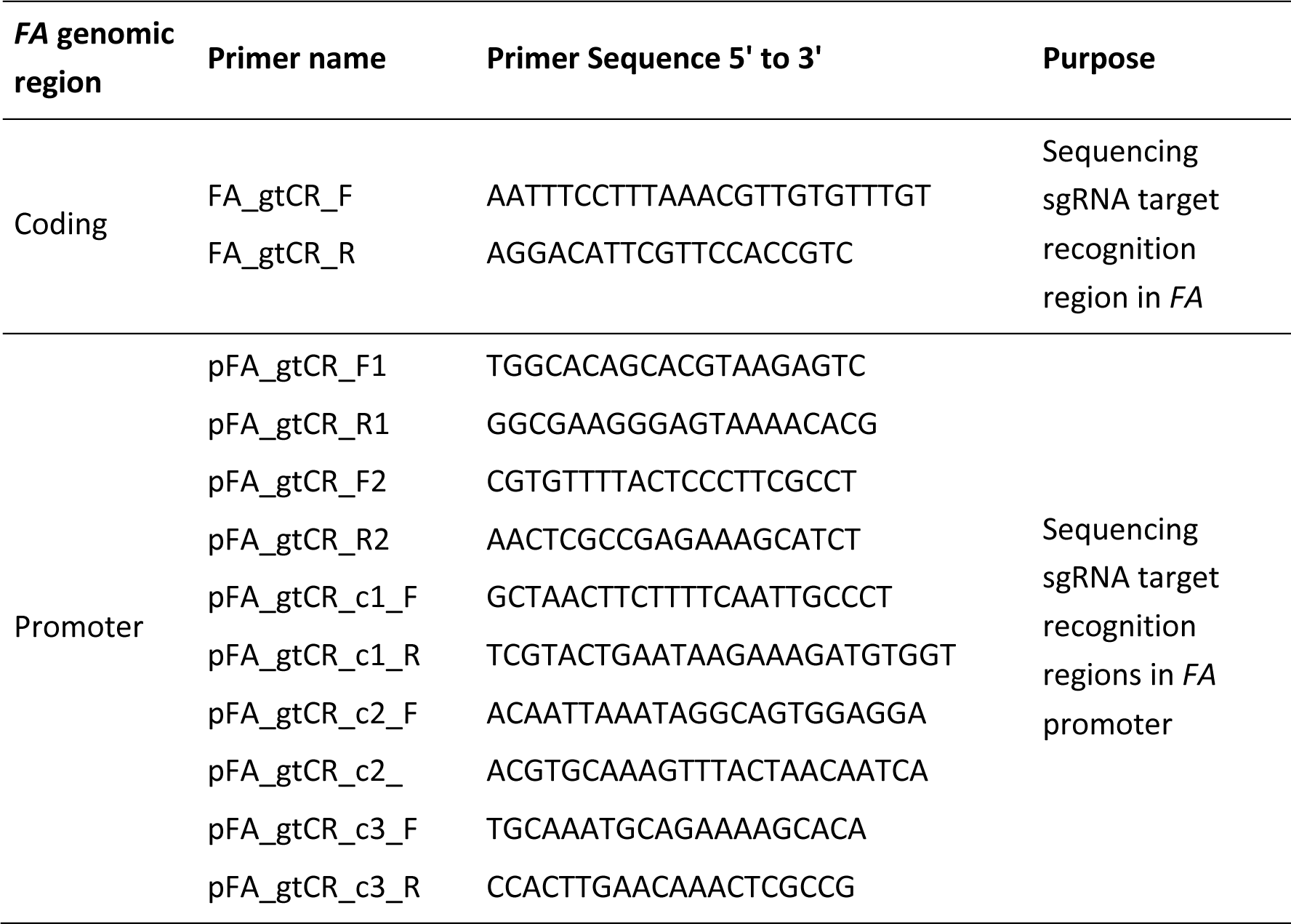
Primer sequences used for sequencing CRISPR/Cas9 edition.

**Supplementary Table S4.**
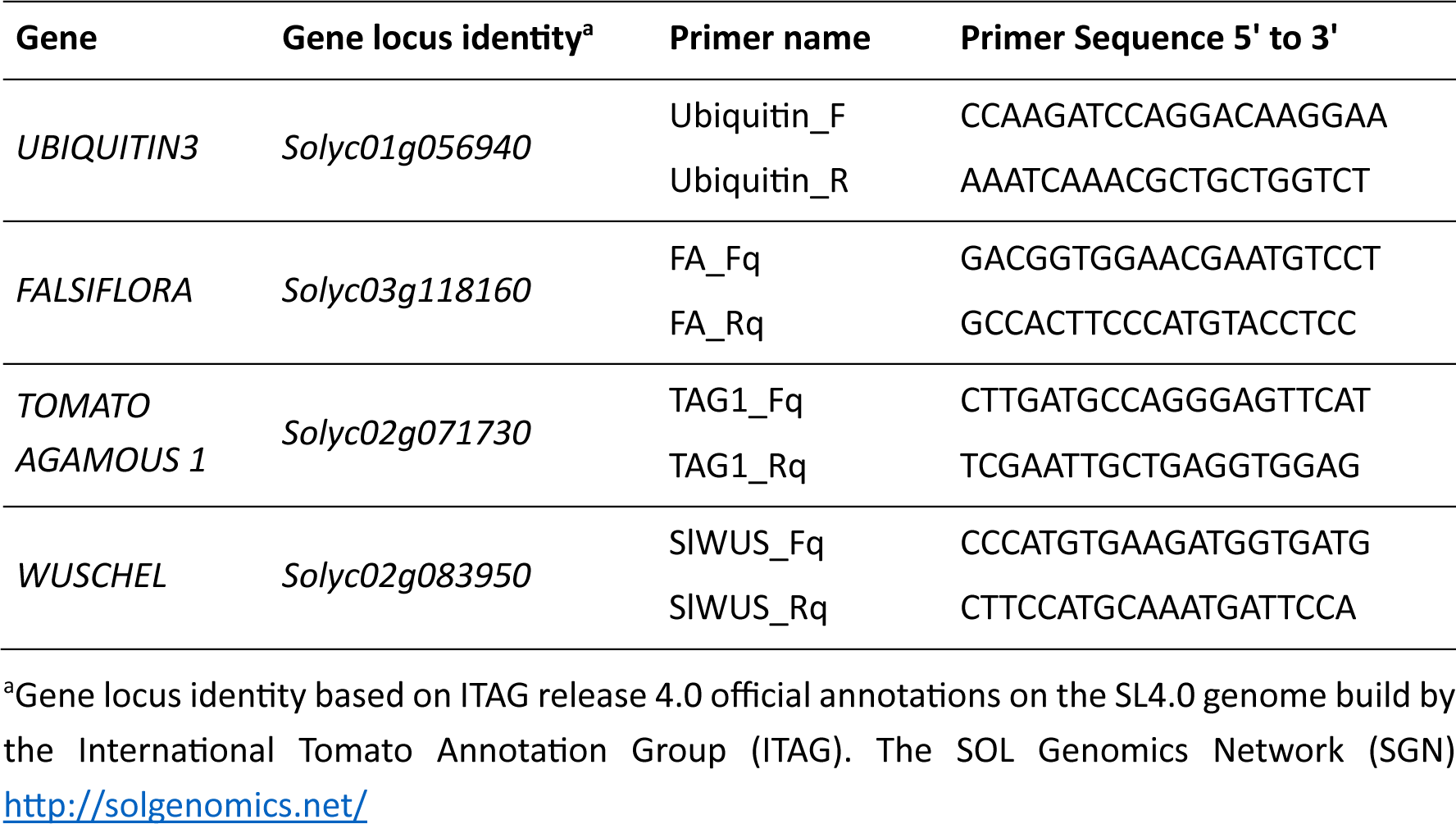
Primer sequences used for qRT-PCR analyses.

